# A role for *Toxoplasma gondii* chloroquine resistance transporter in bradyzoite viability and digestive vacuole maintenance

**DOI:** 10.1101/647669

**Authors:** Geetha Kannan, Manlio Di Cristina, Aric J. Schultz, My-Hang Huynh, Fengrong Wang, Tracey L. Schultz, Matteo Lunghi, Isabelle Coppens, Vern B. Carruthers

## Abstract

*Toxoplasma gondii* is a ubiquitous pathogen that can cause encephalitis, congenital defects, and ocular disease. *T. gondii* has also been implicated as a risk factor for mental illness in humans. The parasite persists in the brain as slow growing bradyzoites contained within intracellular cysts. No treatments exist to eliminate this form of parasite. Although proteolytic degradation within the parasite lysosomal-like vacuolar compartment (VAC) is critical for bradyzoite viability, whether other aspects of the VAC are important for parasite persistence remains unknown. An ortholog of *Plasmodium falciparum* CRT has previously been identified in *T. gondii* (TgCRT). To interrogate the function of TgCRT in chronic stage bradyzoites and its role in persistence, we knocked out TgCRT in a cystogenic strain and assessed VAC size, VAC digestion of host-derived proteins and parasite autophagosomes, and viability of *in vitro* and *in vivo* bradyzoites. We found that whereas parasites deficient in TgCRT exhibit normal digestion within the VAC, they display a markedly distended VAC and their viability is compromised both *in vitro* and *in vivo*. Interestingly, impairing VAC proteolysis in TgCRT deficient bradyzoites restored VAC size, consistent with a role for TgCRT as a transporter of products of digestion from the VAC. In conjunction with earlier studies, our current findings suggest a functional link between TgCRT and VAC proteolysis. This work provides further evidence of a crucial role for the VAC in bradyzoite persistence and a new potential VAC target to abate chronic *Toxoplasma* infection.

**IMPORTANCE:** Individuals chronically infected with the intracellular parasite *Toxoplasma gondii* are at risk of experiencing reactivated disease that can result in progressive loss of vision. No effective treatments exist for chronic toxoplasmosis due in part to a poor understanding of the biology underlying chronic infection and a lack of well validated potential targets. Here we show that a *T. gondii* transporter is functionally linked to protein digestion within the parasite lysosome-like organelle and that this transporter is necessary to sustain chronic infection in culture and in experimentally infected mice. Ablating the transporter results in severe bloating of the lysosome-like organelle. Together with earlier work, this study suggests the parasite’s lysosome-like organelle is vital for parasite survival, thus rendering it a potential target for diminishing infection and reducing the risk of reactivated disease.

## INTRODUCTION

*Toxoplasma gondii* (*T. gondii*) is an opportunistic pathogen that causes encephalitis or debilitating ocular and congenital diseases in humans (1–4). It has also been implicated as a risk factor for schizophrenia and other major mental illnesses (5–8). The parasite progresses through two major life stages during infection of its intermediate hosts: the acute stage characterized by actively replicating tachyzoites and the chronic stage featuring slow growing bradyzoite cysts that persist in muscle and brain tissue (9). While drugs exist against acute stage tachyzoites, currently no treatments are available to combat the chronic stage bradyzoite cysts. The development of new interventions for limiting disease from chronic infection is hindered by a lack of well-validated potential targets and understanding of the biology of *T. gondii* bradyzoites.

One avenue toward this goal is to define the contributions of proteins associated with the parasite Plant-Like Vacuole (PLV)/Vacuolar Compartment (VAC, used hereafter). The *T. gondii* VAC is a lysosome-like organelle that contains a variety of proteases, including those of the cathepsin family (10, 11). It was previously shown that *T. gondii* cathepsin protease L (TgCPL) localizes to the lumen of the VAC where it aids in the digestion of ingested host-derived proteins and parasite autophagosomes (11–13). Diminishing the digestive function of the VAC by either genetic ablation of TgCPL or chemical inhibition of TgCPL with morpholinurea-leucine-homophenylalanine-vinyl phenyl sulphone (LHVS) revealed an critical role for the VAC in parasite viability, particularly in the bradyzoite stage (11, 13, 14).

The *T. gondii* VAC also possesses transmembrane proteins, including an orthologue of the *Plasmodium falciparum* chloroquine resistance transporter (PfCRT) (15). *Arabidopsis thaliana* expresses a homologue of PfCRT as well, which is involved in export of glutathione from plant chloroplasts (16). Similarly, PfCRT has been implicated in the transport of amino acids and peptides out of the digestive vacuole, and is also important for efflux of chloroquine from the malaria digestive vacuole to the parasite cytosol (17). Recent work utilizing yeast demonstrated that *T. gondii* CRT (TgCRT) is also capable of transporting chloroquine (18). Thus, similar to PfCRT, *T. gondii* CRT (TgCRT) might also transport small amino acids and peptides out of the *T. gondii* VAC and into the parasite cytosol. Two studies have revealed that *T. gondii* RH tachyzoites deficient in TgCRT, either by inducible knockdown or complete genetic ablation, exhibit an enlarged VAC (15, 18). In addition, expansion of the VAC in TgCRT deficient tachyzoites is diminished when parasite digestion is impaired by genetic ablation of cathepsin protease B (TgCPB) or chemical inhibition of TgCPL with LHVS (18). Thus, the distended VAC in TgCRT deficient tachyzoites was postulated to be due to increased osmotic pressure from a buildup of digestion products that could not be transported out of the VAC (15, 18). TgCRT deficient tachyzoites also grow more slowly *in vitro* and are compromised in their ability to cause mortality in mice during acute infection, suggesting an inability to transport digested material out of the VAC and into the parasite cytosol has a moderate effect on *T. gondii* tachyzoites (15, 18)

However, the extent to which TgCRT functions as a transporter of digestion products in bradyzoite cysts and thereby contributes to VAC morphology or function, or whether it is necessary for parasite viability during the chronic stage of infection, is unknown. We therefore sought to define the function of TgCRT in bradyzoites and its contribution to bradyzoite survival. To study this, we created a knockout of TgCRT in a cystogenic strain and assessed VAC morphology, *in vitro* and *in vivo* viability, and VAC digestion of host- or parasite-derived material in TgCRT deficient bradyzoites. We show that these bradyzoites exhibit a severely bloated VAC, that TgCRT appears to function downstream of protein digestion within the VAC, and that TgCRT deficiency results in loss of bradyzoite viability.

## RESULTS

### PΔ*crt* parasites exhibit a markedly distended VAC

To examine the role of TgCRT in bradyzoites, we knocked out TgCRT in the cystogenic type II Prugniaud strain (PΔ*crt*) and restored its expression via genetic complementation (PΔ*crt:CRT*) (Fig. S1 & S2). Consistent with TgCRT playing a role in VAC morphology, PΔ*crt* extracellular tachyzoites (Fig. 1A) and bradyzoites isolated from *in vitro* cysts (Fig. 1B) show a larger translucent vacuole associated with the VAC marker TgCPL than the parental and complement strains. The translucent vacuole was also observed within intact *in vitro* TgCRT deficient bradyzoite cysts, as seen by phase contrast (Fig. 1C) and electron microscopy (EM) (Fig. 1D), suggesting VAC enlargement in bradyzoites is not strictly a consequence of being in an extracellular environment. Quantification of EM images reveals a 5-fold enlargement of VAC area in PΔ*crt* bradyzoites as compared with the parental and complement strains (Fig. 1E). These results indicate that TgCRT deficiency in a cystogenic type II strain results in a pronounced enlargement of the VAC in both tachyzoites and bradyzoites.

**Figure 1.**
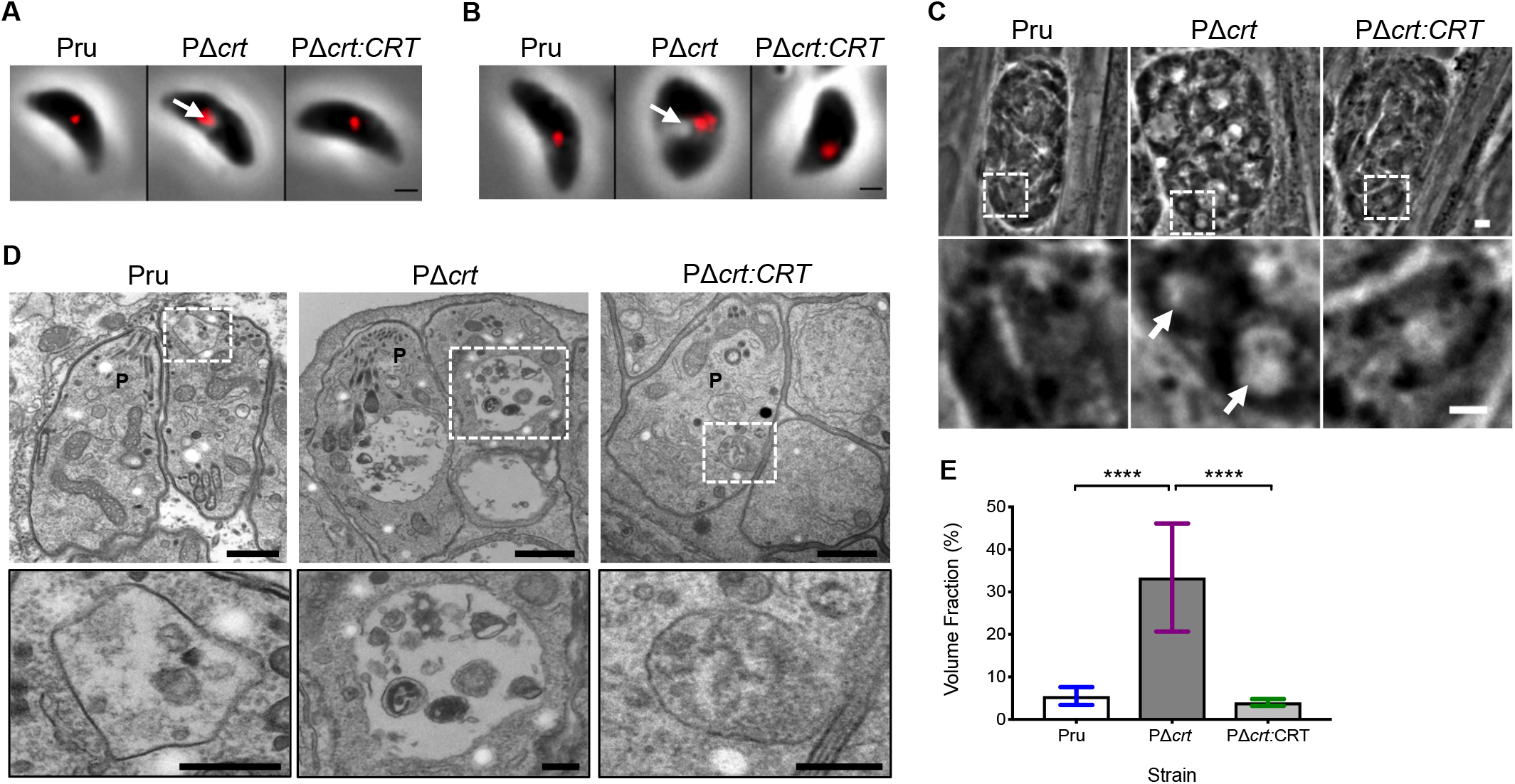
PΔ*crt* tachyzoites and bradyzoites exhibit a distended VAC. A. Extracellular tachyzoites stained for the VAC protease TgCPL (red). Scale bar denotes 1 μM. Arrow denotes distended VAC. B. Extracellular bradyzoites purified from *in vitro* cysts differentiated for 1 week and stained for TgCPL (red). Scale bar denotes 1 μM. Arrow denotes distended VAC. C. Intracellular bradyzoite cysts differentiated *in vitro* for 1 week. Scale bar denotes 10 μM. Arrow denotes distended VAC. D. Electron micrographs of intracellular bradyzoite cysts cultured *in vitro* for 1 week. Images within white boxes were expanded for the insets shown in the second row. Scale bars represent 500 nm for low magnification images and 200 nm for insets. P denotes the parasite. E. Quantification of VAC size from electron micrographs. The following number of VACs were measured for each strain: Pru (13), PΔ*crt* (25), PΔ*crt:CRT* (15). Volume fraction corresponds to the area of the VAC/area of the parasite x 100. Bar is mean +/− SD. One-way ANOVA with Tukey’s multiple comparisons was performed. **** denotes *p*<0.0001.

### TgCRT is required for *in vitro* bradyzoite viability and *in vivo* cyst burden

Previous work established that proteolytic digestion of material in the VAC is necessary for survival of *T. gondii* bradyzoites *in vitro* and *in vivo* (13). Because TgCRT is important for maintaining normal VAC morphology, we reasoned that TgCRT deficiency might compromise bradyzoite viability. We first wanted to address whether the lack of TgCRT affected the rate or efficiency of tachyzoite to bradyzoite conversion and bradyzoite replication. The parasite strains used express GFP under the early bradyzoite LDH2 promoter (19). To assess conversion, we measured the percentage of parasite-containing vacuoles that were greater than 50% positive for GFP or the more mature stage bradyzoite specific marker TgBAG1 over the first 4 days of conversion. For both early- and mature-bradyzoite stage markers analyzed, we found that all strains converted at a similar rate (Fig. S3). In addition, we measured the cyst size as an indicator of bradyzoite replication at 1 and 2 weeks post-conversion and found them to be comparable amongst all strains at both time points (Fig. S4). These findings suggest that TgCRT is not necessary for acute to chronic stage differentiation or replication of chronic stage parasites up to 2 weeks *in vitro*.

We then sought to assess the extent to which TgCRT deficiency affects bradyzoite viability *in vitro*. First we measured the expression of GFP as a proxy of bradyzoite health. It was previously shown that as bradyzoite viability decreases, there is a shift from cysts being uniformly GFP positive to partially positive (mixture of GFP positive and GFP negative) to fully GFP negative (13). Although we found that PΔ*crt* cysts were uniformly GFP positive (Fig. 2A) the intensity of GFP was diminished at 2 weeks, but not 1 week, post-conversion (Fig. 2B), suggesting a temporal decrease in gene expression. Next we more directly evaluated bradyzoite viability using a qPCR/plaque assay (13), which measures the ability of bradyzoites to initiate plaque formation relative to the inoculum (plaques/1000 genomes). We found that PΔ*crt* bradyzoite viability was decreased at 2 weeks, but not 1 week, post-conversion (Fig. 2C), mirroring the findings for GFP intensity. As a decrease in plaques/genomes could be attributed to a deficiency in the ability of PΔ*crt* parasites to form plaques, we conducted a tachyzoite plaque assay that revealed PΔ*crt* tachyzoites have no deficit in the number of plaques formed (Fig. S5). Together these findings indicate a progressive loss of PΔ*crt* bradyzoite viability *in vitro*.

**Figure 2.**
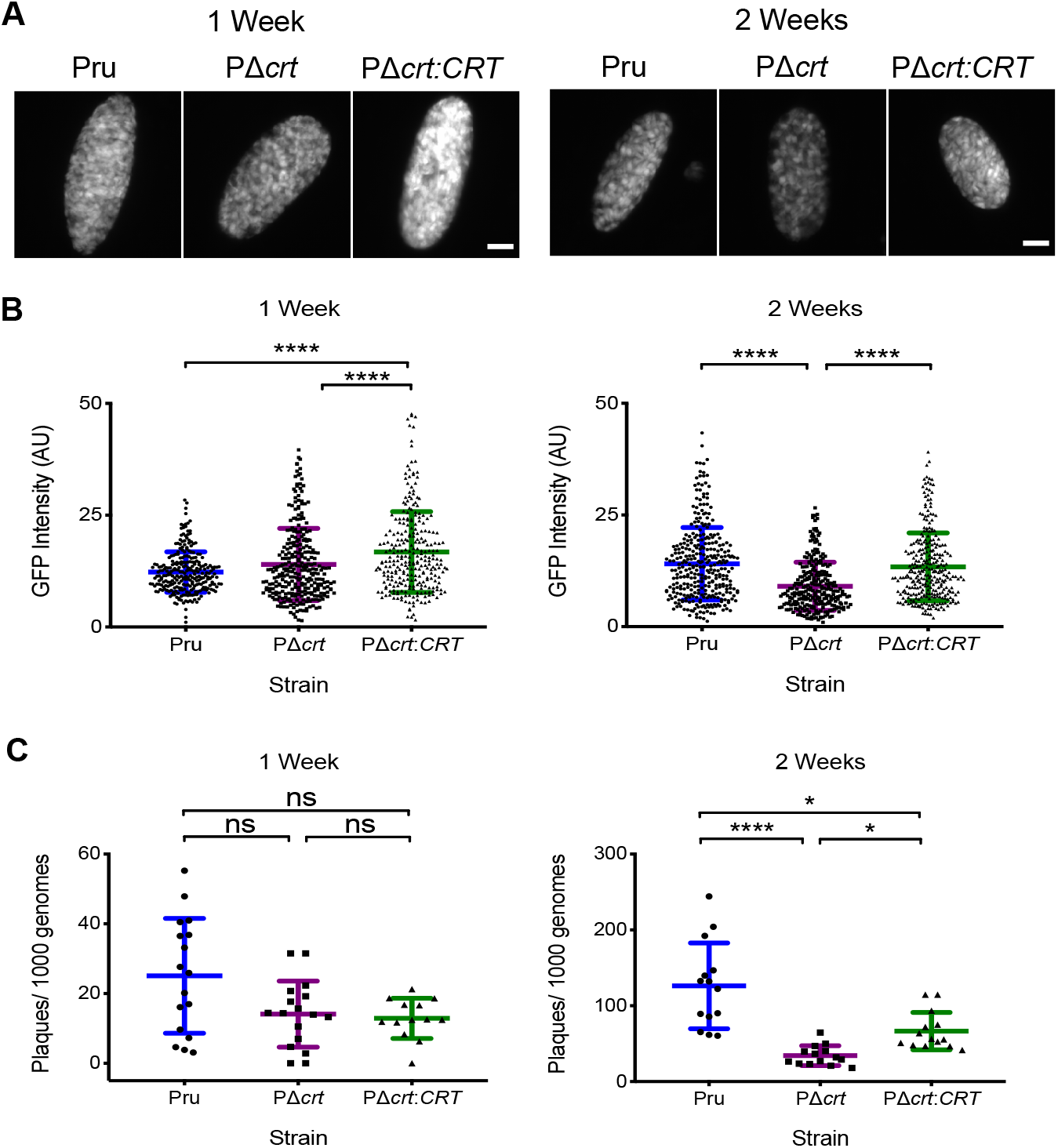
Deletion of CRT affects *in vitro* bradyzoite viability. A. Fluorescent images of bradyzoite cysts expressing GFP under the early bradyzoite LDH2 promoter after 1 and 2 weeks of *in vitro* differentiation. Scale bars denote 10 μM. B. GFP intensity after 1 and 2 weeks of *in vitro* differentiation. Line represents mean +/− S.D. of bradyzoite cysts in 3 independent experiments. The following number of cysts were analyzed for each experiment for week 1: Pru (77, 54, 142), PΔ*crt* (54, 91, 148), PΔ*crt:CRT* (96, 92, 88) and for week 2: Pru (106, 124, 102), PΔ*crt* (56, 131, 94), PΔ*crt:CRT* (89, 107, 102). Kruskal Wallis with Dunn’s multiple comparisons was performed. **** denotes p<0.0001. C. Viability of bradyzoites after 1 and 2 weeks of *in vitro* differentiation based on plaque numbers normalized to qPCR quantification. Line represents mean +/− S.D. of 3-4 technical replicates in 4-5 independent experiments. The following number of technical replicates were analyzed for each experiment for week 1: Pru (3, 3, 3, 4, 4), PΔ*crt* (3, 3, 3, 4, 4), PΔ*crt:CRT* (3, 3, 3, 4) and for week 2: Pru (3, 3, 3, 4, 4), PΔ*crt* (3, 3, 3, 4, 4), PΔ*crt:CRT* (3, 3, 3, 4, 4). Kruskal Wallis with Dunn’s multiple comparisons was performed. **** denotes *p*<0.0001 and * denotes *p*<0.05.

To determine whether deletion of TgCRT affects the chronic infection *in vivo* we infected C57BL/6 mice and enumerated brain cysts at 4 weeks post-infection. Mice inoculated with PΔ*crt* tachyzoites showed a ~10-fold decrease in brain cyst burden compared with those inoculated with the parental or complement strains (Fig. 3A). The reduction in cyst burden was not due to a lack of infection since all mice were seropositive for *T. gondii* IgG, including those in which no cysts were observed (Fig. 3B). However, it is possible that the reduced number of PΔ*crt* brain cysts observed was due to fewer tachyzoites entering the brain during acute infection. To examine this, we used qPCR to measure initial levels of infection in the brain at days 7 and 10 post-infection. Compared to those infected with parental or complement strains, mice infected with PΔ*crt* parasites showed a 2-3 fold lower brain burden, suggesting that the decrease in cyst burden at 5 weeks post-infection is partly attributable to lower initial infection of the brain.

**Figure 3.**
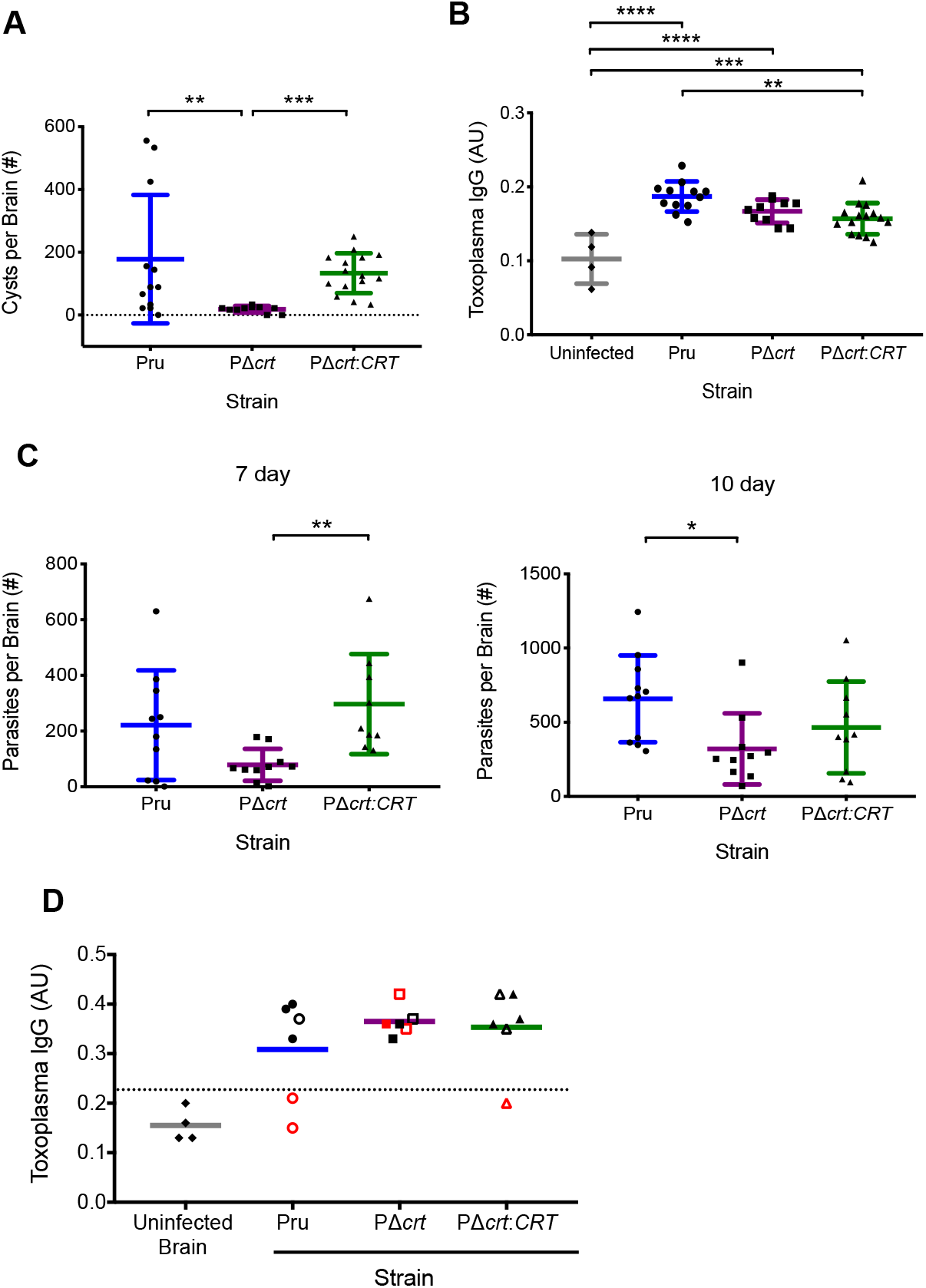
Deletion of TgCRT affects *in vivo* bradyzoite burden. A. Brain cyst burden in mice at 4 weeks post-infection with *T. gondii*. Line represents mean +/− S.D. of mice from 2 independent experiments. The total number of mice analyzed were: Pru (12), PΔ*crt* (10), PΔ*crt:CRT* (15). Kruskal-Wallis with Dunn’s multiple comparisons was performed. *** denotes *p*=0.0002 and ** denotes *p*= 0.0098. B. *T. gondii* IgG of mice infected in panel A. Age and sex-matched uninfected mice used as IgG negative control. One-way ANOVA with Holm-Sidak’s multiple comparisons was performed. **** denotes *p*<0.0001, *** denotes *p*=0.0002, ** denotes *p*=0.002. C. Brain parasite burden at 7 and 10 dpi. Line is mean +/− S.D. of mice from 2 independent experiments. Kruskal-Wallis with Dunn’s multiple comparisons was performed. The following is the number of mice analyzed for 7 and 10 dpi respectively: Pru (10,11), PΔ*crt* (10,10), PΔ*crt:CRT* (9,10). ** denotes *p*=0.005, * denotes *p*=0.017. D. *T. gondii* IgG levels in mice administered residual brain cysts (5 or 30 cysts). Data is from 1 experiment. Line is mean and dotted line is 2 SD above the mean of mice given uninfected brain homogenate. Open symbols denote mice administered 5 parasite cysts and closed symbols denote mice administered 30 parasite cysts. Red symbols denote no parasite growth from brain homogenate. The following is the total number of mice analyzed: Uninfected mice (4), Pru (6), PΔ*crt* (6), PΔ*crt:CRT* (6).

Because we found that *in vitro* TgCRT deficient bradyzoites are less viable, we wanted to examine whether residual *in vivo* PΔ*crt* cysts are infectious. To test this, we inoculated naïve mice with 5 or 30 cysts from the brains of mice chronically infected with Pru, PΔ*crt*, or PΔ*crt:CRT*. Once in the chronic phase, infection of naïve mice was monitored via serology and by determining whether parasites could be cultured from their brain homogenates. To serve as a negative control, 5 naïve mice were inoculated with brain homogenate from an uninfected mouse. All mice inoculated with PΔ*crt* brain cysts were seropositive, indicating that PΔ*crt* cysts contain infectious bradyzoites (Fig. 3D). However, only 50% of the seropositive mice were culture positive. In contrast, while not all mice inoculated with parental or complement brain cysts were seropositive, parasites were cultured from the brains of 100% of the seropositive mice. Taken together, our *in vitro* and *in vivo* data indicate that TgCRT deficient bradyzoites show a decrease, but not absolute loss, of viability.

### Digestion in the VAC of TgCRT deficient tachyzoites and bradyzoites

We next wanted to interrogate whether the decreased viability in TgCRT deficient bradyzoites is possibly due to an impairment of proteolytic digestion in the VAC. Pru strain tachyzoites and bradyzoites deficient in the VAC protease TgCPL (PΔ*cpl*) have a deficit in digestion and reduced bradyzoite viability (13). It was recently suggested that RHΔ*crt* tachyzoites have 25% less TgCPL, but the extent to which this affects VAC digestion was not assessed (18). To probe whether TgCRT deficiency affects VAC digestion in tachyzoites, we utilized a tachyzoite ingestion/digestion assay that permits the detection of ingested and undigested host-derived mCherry within tachyzoites (12). We included PΔ*cpl* as a reference control since these parasites accumulate host-derived mCherry due to a deficiency in VAC proteolytic activity (11, 13, 14). We also created a PΔ*crt*Δ*cpl* double knockout strain by ablating TgCRT in the PΔ*cpl* strain to determine whether a lack of accumulated host-derived material in PΔ*crt* parasites is due to functional digestion or problems in protein delivery to the VAC (Fig. S1). Western blotting confirmed that TgCPL was expressed in all strains except for PΔ*cpl* and PΔ*crt*Δ*cpl* (Fig. 4A). Accumulation of host-derived mCherry was observed in tachyzoites of all strains (Fig. 4B). However, we found that whereas 33% of PΔ*cpl* and 38% PΔ*crt*Δ*cpl* tachyzoites accumulated host-derived mCherry, PΔ*crt* showed only 3% mCherry positive tachyzoites, which is comparable to the parental and complement lines (Fig. 4C). Accumulation of mCherry in PΔ*crt*Δ*cpl* parasites was not significantly different than that of PΔ*cpl*. Taken together, these findings suggest that TgCRT is not required for the delivery or digestion of host-derived protein in the VAC of tachyzoites.

**Figure 4.**
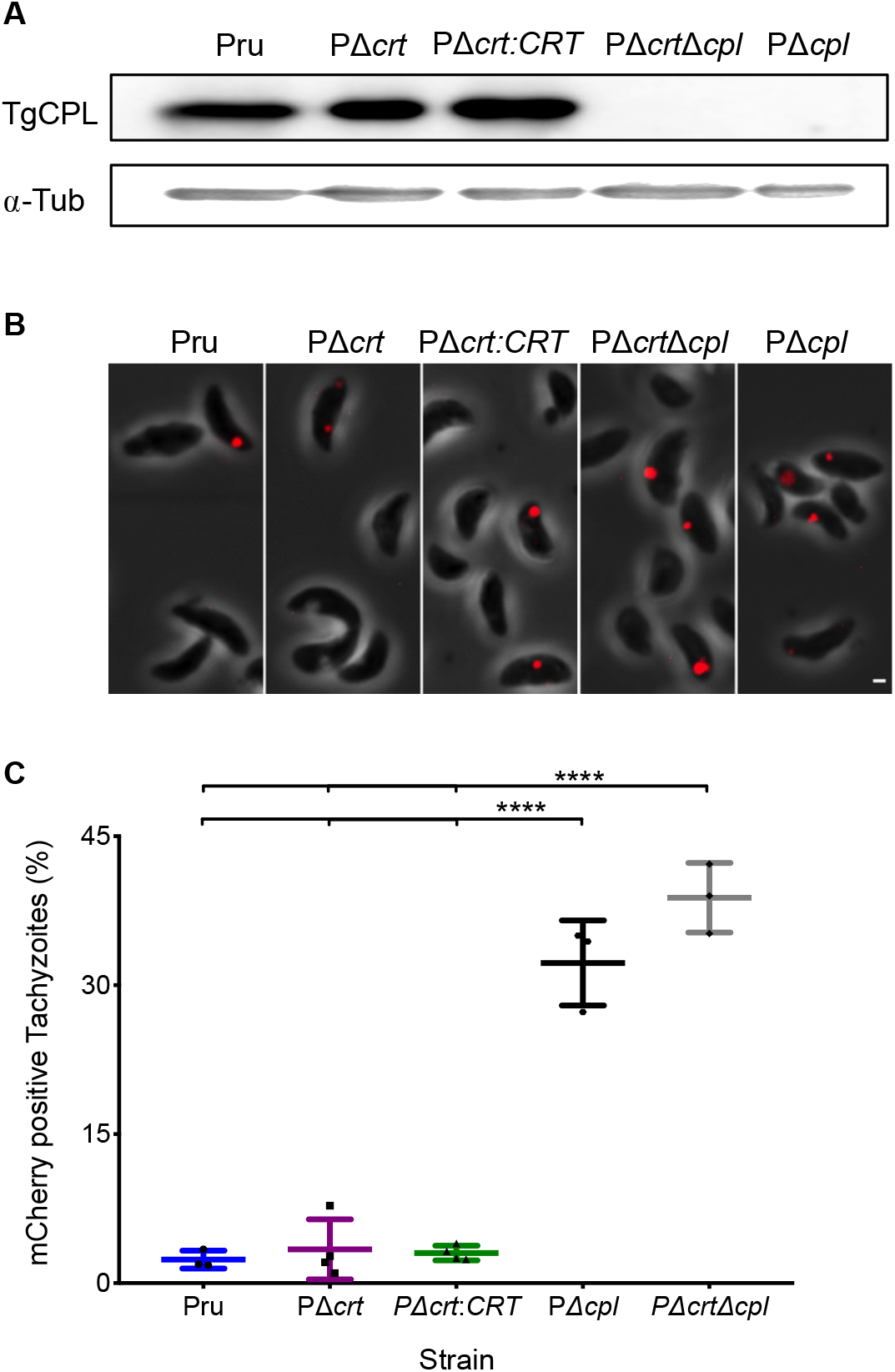
VAC digestive function is not altered in PΔ*crt* tachyzoites. A. Western blot of tachyzoite lysates probed for TgCPL (~30 kDa) and α-Tubulin (~55 kDa) as loading control. B. Representative images of tachyzoites with ingested host-derived mCherry in red. Scale bar denotes 1 μM. C. Tachyzoite ingestion/digestion assay quantification from panel B. Lines represent the mean +/− SD of 3-4 experiments. The following numbers of tachyzoites were enumerated for each experiment: Pru (234, 370, 280), PΔ*crt* (297, 258, 290, 241), PΔ*crt:CRT* (235, 282, 239, 466), PΔ*crt*Δ*cpl* (268, 211, 270), PΔ*cpl* (426. 384, 275). One-way ANOVA with Holm-Sidak’s Multiple Comparisons was performed. **** denotes p<0.0001.

We next wanted to determine whether TgCRT deficiency affects VAC digestion in bradyzoites. Since it has not yet been shown whether bradyzoites are capable of ingesting host cytosolic material akin to tachyzoites, we instead employed a ‘puncta’ assay to initially assess VAC digestion in bradyzoites. This assay is based on a previous report showing that disruption of VAC proteolysis with the TgCPL inhibitor LHVS leads to the accumulation of undigested material in the VAC, which is visible by phase contrast microscopy as dark puncta (13). We found that PΔ*crt* cysts treated with LHVS developed dark puncta and that this corresponded with loss of the translucent VAC (Fig. 5A and B). As expected, there was an increase in dark puncta of parental and complement LHVS treated cysts as well. However, PΔ*crt* cysts contain larger dark puncta in both DMSO and LHVS treated samples than in the parental and complement cysts (Fig. 5B). Also, although PΔ*crt* bradyzoites did not show an increase in the total number of puncta (Fig. 5C), the percentage of total cyst area occupied by puncta was increased with LHVS treatment (Fig. 5D). Together these findings suggest that PΔ*crt* bradyzoites have larger puncta as an indicator of undigested material; however, whether this is a result of moderately impaired proteolytic digestion within the VAC or the intrinsically larger size of PΔ*crt* VAC is unclear.

**Figure 5.**
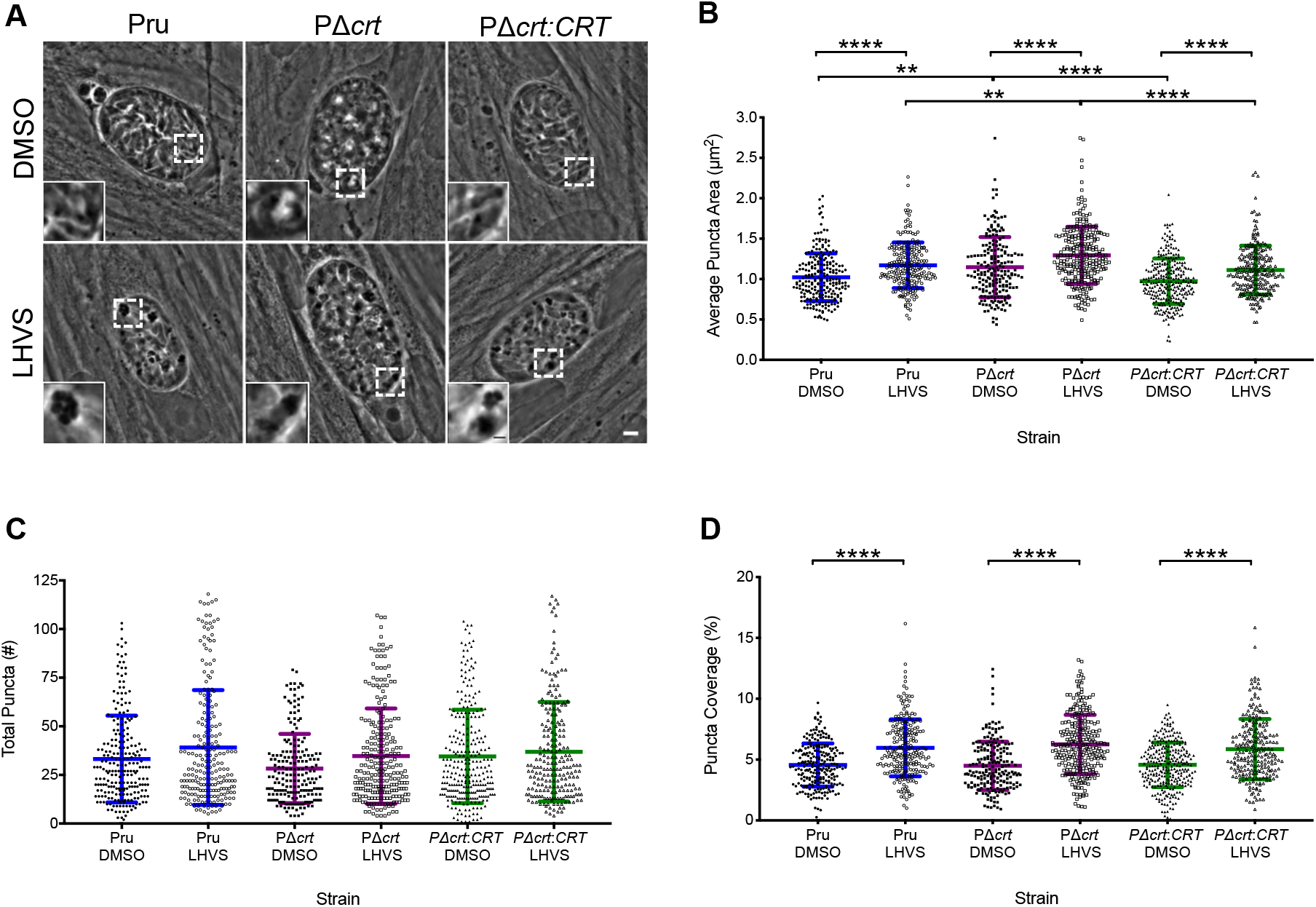
VAC digestive function is not altered in PΔ*crt* bradyzoites. A. Representative images of bradyzoite cysts cultured *in vitro* for 7 days and then treated with DMSO as a vehicle control or 1 μM LHVS for 3 days. Dark puncta are clearly seen in LHVS treated cysts. Scale bar represents 5 μM and scale bar of inset represents 1 μM. B. Measurement of dark puncta area within cysts from 2-3 independent experiments. Lines represent mean +/− S.D. The following number of cysts were analyzed from each experiment: Pru DMSO (65, 68), Pru LHVS(66, 73), PΔ*crt* DMSO(72, 59, 69), PΔ*crt* LHVS(109, 78, 70), PΔ*crt:CRT* DMSO(115, 60, 105), PΔ*crt:CRT* LHVS(77, 56, 94). Kruskal Wallis with Dunn’s multiple comparisons was performed. **** denotes *p*<0.0001, ** denotes *p*<0.01. C. Total puncta number in cysts analyzed in B. Lines represent mean +/− S.D. The following number of cysts were analyzed from each experiment: Pru DMSO (63, 64), Pru LHVS(63, 69), PΔ*crt* DMSO(70, 58, 66), PΔ*crt* LHVS(107, 74, 65), PΔ*crt:CRT* DMSO(112, 58, 106), PΔ*crt:CRT* LHVS(73, 56, 87). Kruskal Wallis with Dunn’s multiple comparisons was performed. D. Percent puncta coverage for each cyst analyzed in B. Lines represent mean +/− S.D. Pru DMSO (65, 68), Pru LHVS(67, 73), PΔ*crt* DMSO(72, 59, 69), PΔ*crt* LHVS(109, 78, 70), PΔ*crt:CRT* DMSO(113, 60, 106), PΔ*crt:CRT* LHVS(77, 56, 94). Kruskal Wallis with Dunn’s multiple comparisons was performed. **** denotes *p*<0.0001.

The dark puncta observed within LHVS-treated bradyzoite cysts have been shown to co-localize with TgCPL and *T. gondii* autophagy-related protein 8 (TgAtg8), suggesting that some of the undigested material found within the bradyzoite VAC is derived from autophagy (13). To interrogate whether TgCRT deficiency affects the production or turnover of parasite autophagosomes, we created a PΔ*crt* strain that ectopically expresses tdTomato-TgAtg8 (Fig. S2), as done previously for Pru (13). Abundance of tdTomato-TgAtg8 in DMSO treated bradyzoites is a function of autophagosomal production and turnover. By contrast, tdTomato-TgAtg8 abundance in LHVS treated bradyzoites is a function of autophagosomal production exclusively since turnover is blocked. Pru- and PΔ*crt* tdTomato-TgAtg8 cysts treated with DMSO or LHVS for 1 or 3 days were assessed for tdTomato-TgAtg8 intensity both within cysts and in isolated bradyzoites. We also measured the total area of tdTomato-TgAtg8 puncta within cysts. For the DMSO control, no significant differences were seen between Pru and PΔ*crt* parasites for tdTomato-TgAtg8 intensity in intact cysts (Fig. 6A & B) or isolated bradyzoites (Fig. 6C), suggesting no change in the balance of autophagosome production and turnover. DMSO treated PΔ*crt* bradyzoites showed a modest, but significant, increase in tdTomato-TgAtg8 puncta size (Fig. 6D), potentially due to tdTomato-TgAtg8 association with the enlarged VAC in such parasites. Upon inhibition of VAC proteolysis with LHVS, tdTomato-TgAtg8 intensity and size increased progressively for both Pru and PΔ*crt* bradyzoites. However, accumulation of tdTomato-TgAtg8 in PΔ*crt* bradyzoites was delayed and somewhat muted compared to Pru. Taken together, these data suggest that the balance of autophagosome production and turnover is unchanged in PΔ*crt*, but that TgCRT deficiency is associated an overall lower rate of autophagosome production.

**Figure 6.**
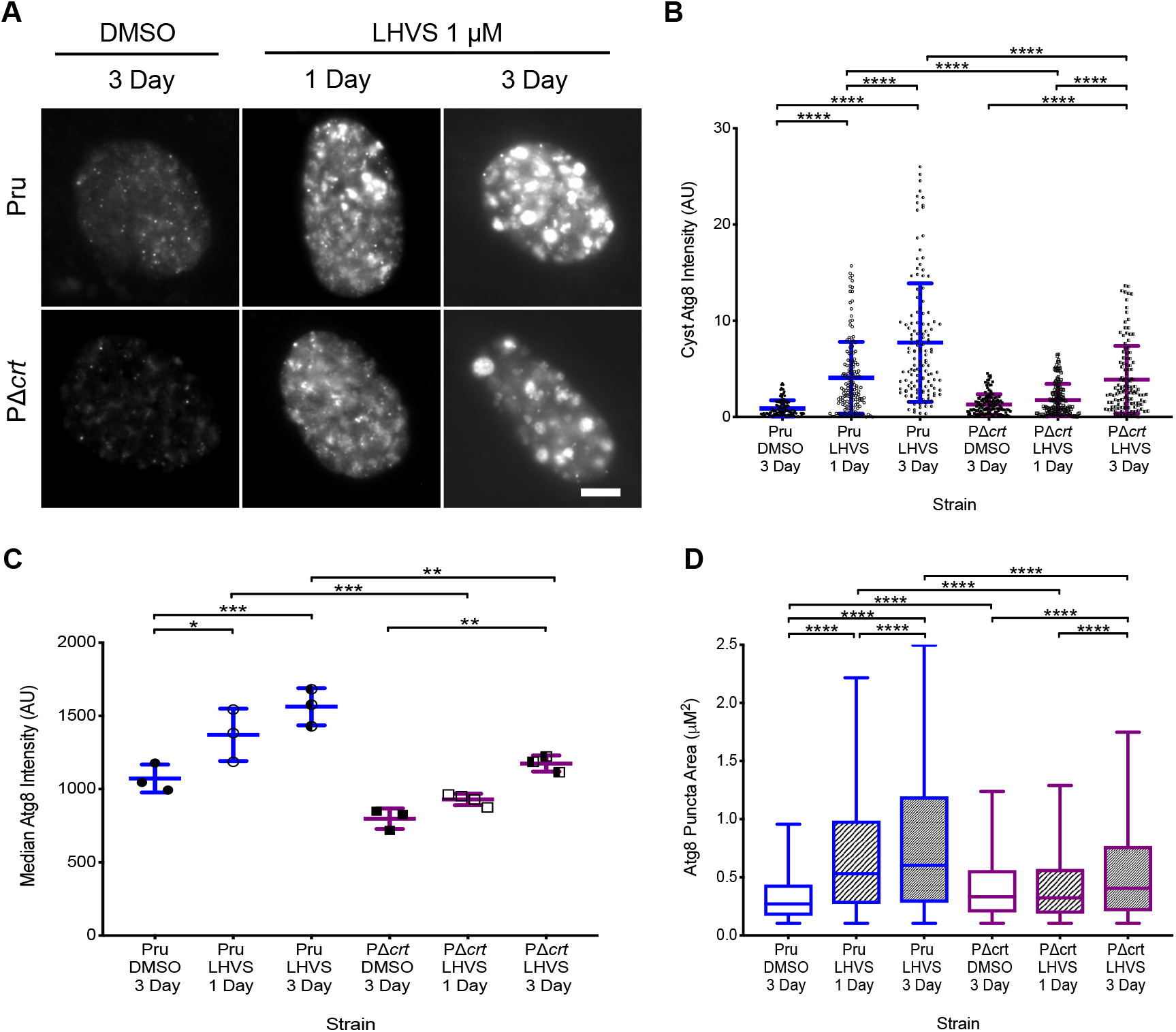
Autophagy in PΔ*crt* bradyzoites. A. Representative images of Pru and PΔ*crt* Atg8-tdTomato expressing strains after 7 days of conversion and treatment with DMSO or 1 μM LHVS for 1 or 3 days. Scale bar represents 10 μM. B. Total tdTomato-TgAtg8 intensity within parasite cysts converted and treated as in A. Line represents mean +/− S.D. from 3-4 independent experiments. The following number of cysts were analyzed in each experiment: Pru DMSO (46, 50, 26), Pru LHVS 1 day (43, 47, 47, 16), Pru LHVS 3 day (45, 60, 31), PΔ*crt* DMSO (48, 54, 30), PΔ*crt* LHVS 1 day (37, 39, 58, 28), PΔ*crt* LHVS 3 day (59, 47, 16). Kruskal Wallis with Dunn’s multiple comparisons was performed. **** denotes *p*<0.0001, ** denotes *p*<0.01. C. Atg8 intensity of bradyzoites analyzed by flow cytometry. Line represents mean +/− S.D. from 3-4 independent experiments. The following number of bradyzoites that were GFP and TdTomato positive were analyzed in each experiment: Pru DMSO (1122, 5330, 1534), Pru LHVS 1 day (493, 3199, 613), Pru LHVS 3 day (1960, 5205, 2043), PΔ*crt* DMSO (620, 1115, 139), PΔ*crt* LHVS 1 day (623, 962, 230, 1355), PΔ*crt* LHVS 3 day (1802, 2641, 337). One-way ANOVA with Sidak’s multiple comparisons was performed. *** denotes *p*<0.001, ** denotes *p*<0.01, * denotes *p*<0.05. D. tdTomato-TgAtg8 puncta size was measured for every puncta in each cyst. Line represents mean +/− S.D. from 3-4 independent experiments. The following number of puncta were analyzed in each experiment: Pru DMSO (364, 290, 242), Pru LHVS 1 day (617, 301, 1826, 1147), Pru LHVS 3 day (722, 697, 1518, 36), PΔ*crt* DMSO (406, 427, 330), PΔ*crt* LHVS 1 day (277, 233, 484, 324), PΔ*crt* LHVS 3 day (692, 402, 633). Kruskal-Wallis with Dunn’s multiple comparisons was performed. **** denotes *p*<0.0001.

### TgCRT transport function is linked to VAC proteolysis

Malaria parasites bearing chloroquine resistance mutations in PfCRT display an enlarged digestive vacuole and they accumulate small peptides derived from hemoglobin (20, 21). This combined with other work showing that recombinant PfCRT transports amino acids, small peptides, and chloroquine (17) suggests that PfCRT functions to transport products of hemoglobin digestion out of the digestive vacuole. More recently, TgCRT was also shown to transport chloroquine upon heterologous expression in yeast (18). It is therefore plausible that TgCRT is also able to transport amino acids and small peptides out of the VAC. If TgCRT plays a similar role and the swelling of the VAC in PΔ*crt* parasites is due to a buildup of TgCRT substrates derived from protein digestion, then reducing the production of digestion products by inhibiting TgCPL should prevent or reverse VAC enlargement.

To test this, we differentiated PΔ*crt* bradyzoites 7 days before adding LHVS for another 2 days under differentiation conditions. This treatment window was chosen because our earlier results showed that 3 days of LHVS treatment results in larger dark and Atg8 puncta areas (Fig. 5B & 6D), whereas a 1 day treatment appeared to have no notable effect on Atg8 intensity (Fig. 5B & C). We reasoned that with 2 days of treatment, we should begin seeing an effect of LHVS treatment on VAC size prior to excessive accumulation of undigested protein. Although some enlarged VACs were apparent in LHVS treated PΔ*crt* bradyzoites (Fig. 7A), quantification revealed a significant restoration of VAC size upon LHVS treatment (Fig. 7B). Also, undigested material accumulated within the VAC of PΔ*crt* bradyzoites treated with LHVS, suggesting that TgCPL is active in PΔ*crt* bradyzoites.

**Figure 7.**
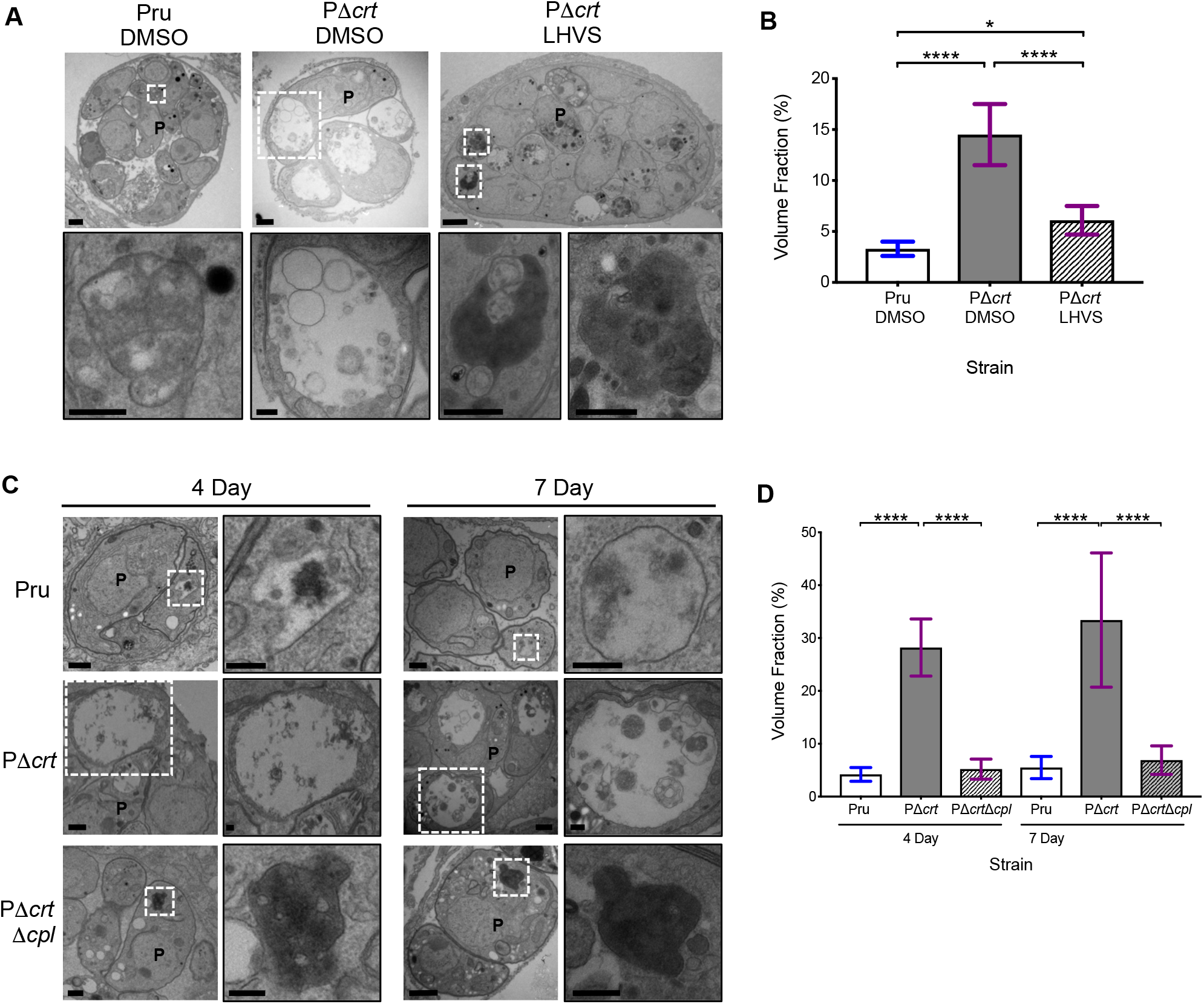
VAC digestion disruption through CPL modulation affects PΔ*crt* bradyzoite VAC size and parasite health. A. Electron microscopy of *in vitro* bradyzoite cysts converted for 7 days and then treated with DMSO and 1 μM LHVS for 2 days. Scale bars represent 500 nm. B. Quantification of VACs in panel A. Bars represent mean +/− S.D. The following numbers of VACs were measured for each strain: Pru (18), PΔ*crt* DMSO (49), PΔ*crt* LHVS (13). One-way ANOVA with Tukey’s multiple comparisons was performed. **** denotes *p*<0.0001, * denotes *p*<0.05. C. Representative electron micrograph images of *in vitro* bradyzoite cysts converted for 4 day and 7 days. Scale bars represent 500 nm. P denotes parasite. D. Quantification of VACs in panel C. Bars represent mean +/− S.D. The following number of VACs were measured for each strain: 4 day Pru (16), PΔ*crt* (17), PΔ*crt*Δ*cpl* (17). 7 day Pru (13), PΔ*crt* (25), PΔ*crt*Δ*cpl* (35). One-way ANOVA with Tukey’s multiple comparisons was performed. **** denotes *p*<0.0001. 7 day Pru and PΔ*crt* data was also used in Figure 1.

To validate a link between TgCRT transport function and VAC proteolysis, we compared the size and appearance of the VAC in PΔ*crt* bradyzoites with that of Pru or PΔ*crt*Δ*cpl* parasites. We found that after 4 or 7 days of conversion to bradyzoite cysts, PΔ*crt*Δ*cpl* bradyzoites have visually smaller VACs full of electron-dense, undigested material compared to the markedly enlarged, more electron-lucent VACs of PΔ*crt* bradyzoites (Fig. 7C). Quantification revealed VAC size of PΔ*crt*Δ*cpl* strains to be significantly smaller than PΔ*crt* VACs at both time points (Fig. 7D). These findings indicate that by genetically limiting proteolysis in the VAC, the gross enlargement of the VAC observed in PΔ*crt* bradyzoites is prevented. In addition, whereas approximately 20% PΔ*crt* cysts were dead or dying at both 4 and 7 days post-conversion, 75% of PΔ*crt*Δ*cpl* cysts were degenerate at 4 days and 100% were degenerate at 7 days post-conversion. Thus, parasite lacking both TgCRT and TgCPL appear to be more severely compromised than those lacking TgCRT alone. Taken together, our findings suggest a link between TgCRT and protein digestion in a manner that is consistent with TgCRT acting as an exporter of degradation products generated by VAC proteases in bradyzoites.

## DISCUSSION

Herein we show that TgCRT is necessary for maintaining the size of the VAC and the viability of *T. gondii* bradyzoites, possibly by functioning as a transporter of digested material from the VAC to the parasite cytosol. Together with other recent studies reporting that VAC protein digestion is crucial for bradyzoite viability (13), our findings point toward the VAC as an important organelle for *T. gondii* bradyzoite persistence and uncover TgCRT as a potential target for chronic *T. gondii* infection.

Our finding that deletion of TgCRT in a type II strain (PΔ*crt*) resulted in enlargement of the VAC is in line with previous studies that have knocked down (15) or knocked out (18) TgCRT in a type I strain (RH). We also show that this enlarged VAC phenotype is consistent across life stages and that it appears to be especially prominent in bradyzoites. Our EM measurements suggest that the VAC occupies one third of the cytoplasm of PΔ*crt* bradyzoites, thus becoming easily visible by phase contract microscopy in many parasites. VAC enlargement was fully reversed upon re-expression of TgCRT, firmly establishing that TgCRT expression is necessary to maintain normal VAC morphology.

The underlying basis for enlargement of the VAC in TgCRT deficient parasites is unknown, but may be linked to endolysosomal system dynamics and the transporter function of TgCRT. The VAC is a dynamic organelle that undergoes rounds of fission to form smaller structures late in the cell cycle before fusing to form typically a single compartment in G1 phase (10, 22). The VAC probably also communicates via fusion and fission with the parasite endosome-like compartments (ELCs), based on partial colocalization of VAC and ELC markers in intracellular parasites (10, 18, 22). Interestingly, it was recently reported that replicating PΔ*crt* parasites maintain a single VAC that overlaps substantially with ELC markers (18). These findings imply that defects in VAC fragmentation and fission of the VAC from the ELCs result in sustaining a hybrid VAC/ELC compartment in parasites lacking TgCRT. Thus, contributions of membrane from both the VAC and ELCs could account for enlargement of the VAC in PΔ*crt* parasites. Although it is possible that TgCRT directly participates in vesicular fission, no evidence of this currently exists. On the other hand, it appears more likely that swelling of the VAC in TgCRT deficient parasites is related to TgCRT transport function. If, akin to PfCRT, TgCRT exports proteolytic digestion products from the VAC, accumulation of such products in the VAC of PΔ*crt* parasites could increase osmotic pressure within the organelle due to the influx of water through a VAC-localized aquaporin (22). Whether a build-up of osmotic pressure is a driver of VAC size in TgCRT deficient parasites and is thereby responsible for defective VAC fragmentation and VAC/ELC resolution awaits further study.

A knockout of *Plasmodium* CRT has not been reported, presumably because of it having an essential function. Nevertheless, chloroquine resistant strains bearing mutations in PfCRT also exhibit an enlarged digestive vacuole. Studies with recombinant PfCRT suggested that chloroquine resistant alleles tend to have lower transport activity for a model substrate (tetraethyl ammonium), but higher transport activity for chloroquine (17). Chloroquine resistant strains also accumulate small peptides derived from digestion of hemoglobin (20, 21). Thus, the enlarged digestive vacuole of chloroquine resistant strains is potentially due to a partial loss of PfCRT native transport function. That PfCRT is essential whereas TgCRT is dispensable likely reflects the crucial role of the malaria digestive vacuole in detoxification of heme liberated from hemoglobin digestion during replication within erythrocytes.

Consistent with an important role for TgCRT in chronic infection, we observed a ~5-fold loss of viability for PΔ*crt* bradyzoites *in vitro*. Loss of viability appears to increase with time of differentiation, suggesting a progressively important role for TgCRT in chronic infection. We also noted a 10-fold decrease in PΔ*crt* brain cysts in mice. This decrease is likely a composite of effects occurring during the acute stage and the chronic stage. The trend toward lower initial infection of the brain observed for PΔ*crt* parasites is in agreement with the decreased virulence reported during acute infection of RHΔ*crt* parasites (18). However, the lower initial infection of the brain does not appear to fully account for the striking decrease in PΔ*crt* brain cysts. Additional loss of PΔ*crt* cysts during the chronic infection of mice is consistent with our *in vitro* viability findings. Nevertheless, we found that cysts recovered from the brains of PΔ*crt* infected mice contained infectious bradyzoites capable of establishing infection of naïve mice. Our observation of lower PΔ*crt* cultivation efficiency from the brains of infected naïve mice is further evidence of a decreased brain burden and/or viability. Thus, whereas TgCRT is not absolutely required for *T. gondii* persistence, it nonetheless strongly influences the course and burden of chronic infection.

Proteolysis within the VAC is necessary for sustaining bradyzoite viability *in vitro* and *in vivo* (13). Genetically or chemically disrupting TgCPL activity results in a loss of bradyzoite viability that is associated with accumulation of undigested material co-localizing with Atg8, a marker of autophagosomes. *P. falciparum* parasites administered protease inhibitors that target digestive vacuole proteinases also accumulate undigested material, in this case hemoglobin derived from the infected erythrocyte (20). However, the electron-lucent VACs observed in PΔ*crt* parasites, along with the tachyzoite ingestion assay, dark puncta measurements, and Atg8 accumulation data suggests that digestion in PΔ*crt* tachyzoites and bradyzoites is largely normal despite the striking morphological changes to the organelle. It was suggested that RHΔ*crt* tachyzoites reduce the expression of several VAC proteases to decrease production of TgCRT substrates generated by VAC proteolysis, thereby easing osmotic pressure (18). If VAC proteolysis is similarly reduced in PΔ*crt* parasites, this does not appear to affect the digestion of host-derived protein in tachyzoites via the ingestion pathway or parasite-derived material delivered through autophagy.

Consistent with TgCRT functioning as a transporter downstream of VAC proteolysis, we found that treating PΔ*crt* bradyzoites with LHVS restored VAC size prior to subsequent accumulation of undigested material. This was observed in the EM images with 2 days of LHVS treatment, where VACs are smaller and a buildup of undigested material is beginning to show. We also found that PΔ*crt*Δ*cpl* double knockout bradyzoites have a normal sized VAC, confirming that protein digestion in the VAC is required for expansion of the VAC in TgCRT deficient parasites. The accumulation of undigested material in LHVS treated PΔ*crt* and PΔ*crt*Δ*cpl* is consistent with delivery of proteolytic substrates to the VAC TgCRT deficient bradyzoites. Nevertheless, we noted a delay in the accumulation of the autophagic marker TgAtg8 in PΔ*crt* bradyzoites after blocking TgCPL activity with LHVS, suggesting a decrease in the production of autophagosomes. Whether this is a result of a feedback loop to reduce delivery of substrates to the VAC akin to the down-regulation of proteases in TgCRT deficient tachyzoites (18) or a due to a general decline in the health of PΔ*crt* bradyzoites remains unclear. It should also be noted that although we were unable to introduce tdTomato-TgAtg8 into PΔ*crt:CRT* parasites due to a lack of available selectable markers, all of the other phenotypes measured in PΔ*crt* parasites were restored upon genetic complementation.

Previous work in TgCPL together with the current findings for TgCRT is consistent with a central role for the VAC in *T. gondii* persistence. Parasites deficient in TgCPL and TgCRT appear to be especially compromised, which is consistent with a functional link between these VAC components. Additional studies aimed at targeting these proteins and identifying new components of the VAC are needed to realize the potential of compromising this organelle for therapeutic gain.

## MATERIALS AND METHODS

### Host cell and parasite cultures

Human foreskin fibroblasts (HFFs) were grown in Dulbecco’s Modified Eagle Medium (DMEM, Gibco) containing 10% cosmic calf (Gibco), 50 μg/ml penicillin-streptomycin, 2 mM L-glutamine, and 10 mM HEPES. *T. gondii* strains used in this study were derived from PruS/Luc strain (13) maintained *in vitro* by serial passage on human foreskin fibroblast (HFF) monolayers as previously described (23).

### VAC staining

Egressed tachyzoites from HFFs were filter-purified and pelleted at 1500x*g* for 10 min. Parasites were settled on Cell-Tak™ (Fisher Scientific) coated slides for 30 min, fixed in 4% formaldehyde, and stained for with RbαTgCPL (1:500) and GtαRb 594 secondary (1:1000).

### *In vitro* conversion

Tachyzoites were converted to bradyzoite cysts *in vitro* using previously published methods (24). In brief, tachyzoites were allowed to invade HFFs overnight under standard growing conditions. Infected cells were then grown in alkaline media (RPMI 1540 w/o NaHCO_3_, 50 mM HEPES, 3% FBS, Pen/Strep, pH 8.2) in an incubator without CO_2_, with media changed every day until samples were processed.

### Transmission electron microscopy

For ultrastructural observations of infected cells by thin-section, samples were fixed in 2.5% glutaraldehyde in 0.1 mM sodium cacodylate and processed as described (25). Ultra-thin sections of infected cells were stained with osmium tetraoxide before examination with Philips CM120 EM (Eindhoven, Netherlands) under 80 kV.

### qPCR/plaque assay for bradyzoite viability

*In vitro* bradyzoite viability was assessed by plaque assays normalized to qPCR as previously described (13). Briefly, tachyzoites were converted to bradyzoite cysts for 7 and 14 days as described above. At these time points bradyzoites were harvested using pepsin treatment and added to HFF monolayers for 10 days, after which time plaques were counted. Genomic DNA was extracted from an aliquot of samples using the Qiagen Blood and Tissue Kit and SYBR Green qPCR performed using the primer pairs listed in Table S1 and the following reaction conditions: 98°C, 2’ [98°C, 5”; 68°C, 30”, 72°C, 45”] x 45 cycles.

### GFP intensity

After 1 and 2 weeks of tachyzoite conversion, as described above, *in vitro* cysts were fixed and stained with biotinylated dolichos (primary; 1:400; Vector Laboratories) and Streptavidin Alexa350 (secondary; 1:1000; Life Technologies). Image J was used to select dolichos stained cysts and quantify the amount of GFP coverage and intensity within the cyst. The dolichos signal was used to create a mask for further analysis by auto thresholding with the Li method (26). Analysis under these masks was redirected to the GFP channel, where particles between 130 – 2300 μm^2^ and 0.30-1.00 circularity were analyzed.

### tdTomato-Atg8 intensity and size

After 1 week of tachyzoite conversion as described above, *in vitro* cysts were fixed and stained with biotinylated dolichos (primary; 1:400; Vector Laboratories) and Streptavidin Alexa350 (secondary; 1:1000; Life Technologies). ImageJ was used to select dolichos stained cysts and quantify the total intensity of tdTomato-Atg8 and the tdTomato-Atg8 puncta size within each cyst. Dolichos positive structures between 200-2000 μM^2^ with a circularity of 0.40-1.00 were identified using the Minimum method of auto-thresholding. The resulting binary images were used to create masks under which Atg8 puncta were further analyzed. Td-Tomato Atg8 puncta were analyzed as being between 0.2-1.50 μM^2^ with a circularity of 0.40-1.00 and were identified with the Phansalkar method of auto local thresholding with a radius of 15.

### Tachyzoite plaque assay

Intracellular tachyzoites were harvested following standard procedures, counted, and added to HFF monolayers in triplicate to quadruplicate wells. Parasites were left undisturbed for 10 days, after which time plaques were counted.

### Mouse seropositivity

*Toxoplasma* IgG was measured using enzyme linked immunosorbent assay (ELISA) to determine infectivity. In brief, *Toxoplasma* lysate was made from freshly lysed Pru tachyzoites that was sonicated in 1 ug/mL Leupeptin, 1 ug/mL E64, TPCK, and 10 ug/mL A-PMSF. Plates were coated with 10 ng of antigen in coating buffer (Na_2_CO_3_, NaHCO_3_, pH 9.6) overnight, blocked in 3% gelatin/PBS-T, serum was added in a 1:25 dilution in 1% gelatin/PBS-T and incubated for 1 hr at RT. Secondary HRP-conjugated GtαMs (1:1000) was added for 1 hr. Substrate was added for color development, which was stopped with H_2_SO_4_. Absorbance was read at 400 nm.

### Tachyzoite ingestion assay

Tachyzoite digestion was determined using the ingestion assay as previously described (12). In brief, inducible mCherry Chinese hamster ovary (CHO) cells were plated and induced with 2 μg/mL of doxycycline for 5 days. Tachyzoites were harvested from HFF cells and allowed to invade induced-CHO cells for 4 hrs. Tachyzoites were then mechanically lysed out of host cells, purified, treated with pronase and saponin, and imaged on Cell-Tak™ (Fisher Scientific) coated slides. Samples were coded at the time of initial harvesting. For each biological replicate, more than 200 tachyzoites of each genotype were analyzed for host-derived mCherry accumulation within parasites.

### Western blotting

Tachyzoite lysates were prepared from purified parasites with the addition of boiled 1x sample buffer, and lysate from 3×10^5^ tachyzoites/10 μL sample buffer was loaded onto 10% SDS polyacrylamide gels. Blots were probed with antibody to TgCPL (Rb; 1:300) (10) and α-tubulin (Ms; 1:1000; Developmental Studies Hybridoma Bank, University of Iowa) for the loading control.

### Puncta measurements in LHVS treated parasites

Tachyzoites were converted to bradyzoite cysts as described above. After 7 days of conversion, parasites were treated with 1 μM LHVS or DMSO (control) every day for 3 days. Cells were then fixed and stained with biotinylated dolichos lectin (primary; 1:400; Vector Laboratories) and Streptavidin Alexa350 (secondary; 1:1000; Life Technologies). Image J was used to select dolichos stained cysts and quantify the number and size of puncta within the cyst. Images were automatically thresholded using the MaxEntropy method to create a binary image (27). Noise was reduced by opening the image with 6 iterations of one pixel. Masks were created by using the Analyze Particle function, with objects between 130 – 1900 μm^2^, and a circularity of 0.30-1.00 begin called a cyst. Under these masks, dark puncta were analyzed in the following way: phase images were Guassian blurred with a sigma of 2, and then auto-local thresholding was performed using the Phansalkar method (28) with a radius of 5 pixels. Objects with an area of 0.20 – 6.00 μm^2^ and a circularity of 0.50 – 1.00 were analyzed as dark puncta.

### *In vitro* differentiation kinetics

Tachyzoites were converted to bradyzoite cysts as described above. Parasites were fixed at 1, 2, 3, and 4 days post-conversion and stained for BAG1 (RbαBAG1, 1:1000), a late marker for bradyzoites. These parasites express GFP under the LDH2 promoter, an early marker of bradyzoites. Image J was used to analyze the BAG1 and GFP coverage of each vacuole. Vacuoles were manually identified using phase images by drawing an ROI with the freehand tool. The ROIs were then applied to other channels for analysis as follows. The GFP and Texas Red channels were auto-thresholded using optimal thresholding methods for each day of conversion. The non-thresholded and thresholded ROIs were measured for pixel intensity and used to determine overall and percent intensity for GFP and Texas Red. Vacuoles with over 50% coverage were designated as being cysts, and the total percentage of GFP and BAG1 positive cysts was calculated independently.

### In *vivo* cyst burden

C57BL/6J female mice (7-8 wks old, Jackson Laboratories, Bar Harbor, ME) were used in this study. Mice were injected intra-peritoneum (i.p.) with purified 10^5^ tachyzoites of either PruS/Luc (Pru), PruΔ*crt*(PΔ*crt*), or PruΔ*crt:CRT* (PΔ*crt:CRT*) in 200 μL of 1x phosphate-buffered saline (PBS). At 4 weeks post-infection (wpi) mice were sacrificed following university-approved protocols. Brains were harvested and homogenized in 1 mL ice-cold PBS via syringing through a 20-gauge needle. Mice were coded and cysts were enumerated in 90 μL of brain homogenate (9% of the brain) and the total brain cyst number calculated. Cyst burden data were pooled from 2 independent experiments.

### *In vivo* parasite burden kinetics

The same inoculation conditions as described for *in vivo* cyst burden was used. At 7 and 10 days post-infection (dpi) mice were sacrificed and brains harvested. Brains were homogenized in ice-cold PBS to have 50 ng homogenate/μL PBS. gDNA was extracted from 50 μL of homogenate using the DNeasy Blood and Tissue Kit (Qiagen). qPCR was performed in triplicate for each sample with the following cycling conditions: 90°C, 2’; [98°C, 10”; 56°C, 20”; 72°C,20"]x45 using SSO Advanced SYBR Green Supermix (BioRad), and 300 nM Tox9 and 11 primers listed in Table S1. *T. gondii* standards of specified parasite numbers (1 – 10^5^genomes/μL) were used to quantify parasite brain burden.

### *In vivo* cyst viability

To determine the viability of *T. gondii* cysts procured from the *in vivo* cyst burden experiment, 5 and 30 brain cysts of Pru, PΔ*crt*, or PΔ*crt:CRT*were injected i.p. into C57BL/6J female mice (7-8 weeks old). Mice inoculated with an equivalent amount of uninfected mouse brain homogenate were used as a negative control for infection. At 3 wpi mice were coded and sacrificed. Serum and brain was collected as described above for the *in vivo* cyst burden. Half of each brain homogenate was added to confluent HFF cells and monitored for parasite growth for 4.5 weeks.

### Flow cytometry

Parasites were fixed with 4% formaldehyde for 15 min at room temperature, washed one time with PBS and resuspended in PBS for analysis on a LSR Fortessa flow cytometer (BD Biosciences, San Jose, CA, USA) with BD FACSDiVa™ Software (BD Biosciences). Data were analyzed with FlowJo (BD Biosciences) using the following gating: FITC-positive parasites were characterized as bradyzoites; then, the amount of tdTomato-ATG8 in bradyzoites was determined by the 561 nm signal.

### Statistics

Data was analyzed using GraphPad prism. For each data set, outliers were identified and removed using ROUT with Q=0.1%. Data was then tested for normality and equal variance. If passed, One-way ANOVA with Tukey’s multiple comparisons was performed. If failed, Mann-Whitney U test or Kruskal-Wallis with Dunn’s multiple comparison was performed.

## ACKNOWLEDGEMENTS

This work was support by National Institutes of health grants R01AI060767 (to I.C.) and R01AI120627 (to V.B.C. and M.D.C.) and a grant from the Stanley Medical Research Institute (to V.B.C.).

We thank the excellent technical staff of the Electron Microscopy Core Facility at the Johns Hopkins University School of Medicine Microscopy Facility. We also thank Professor Carla Emiliani for helpful discussions and support. We appreciate the technical assistance of Drs. Abimbola Kolawole and Carmen Mirabelli with flow cytometry.

## SUPPLEMENTARY MATERIAL LEGENDS

**Table S1.**
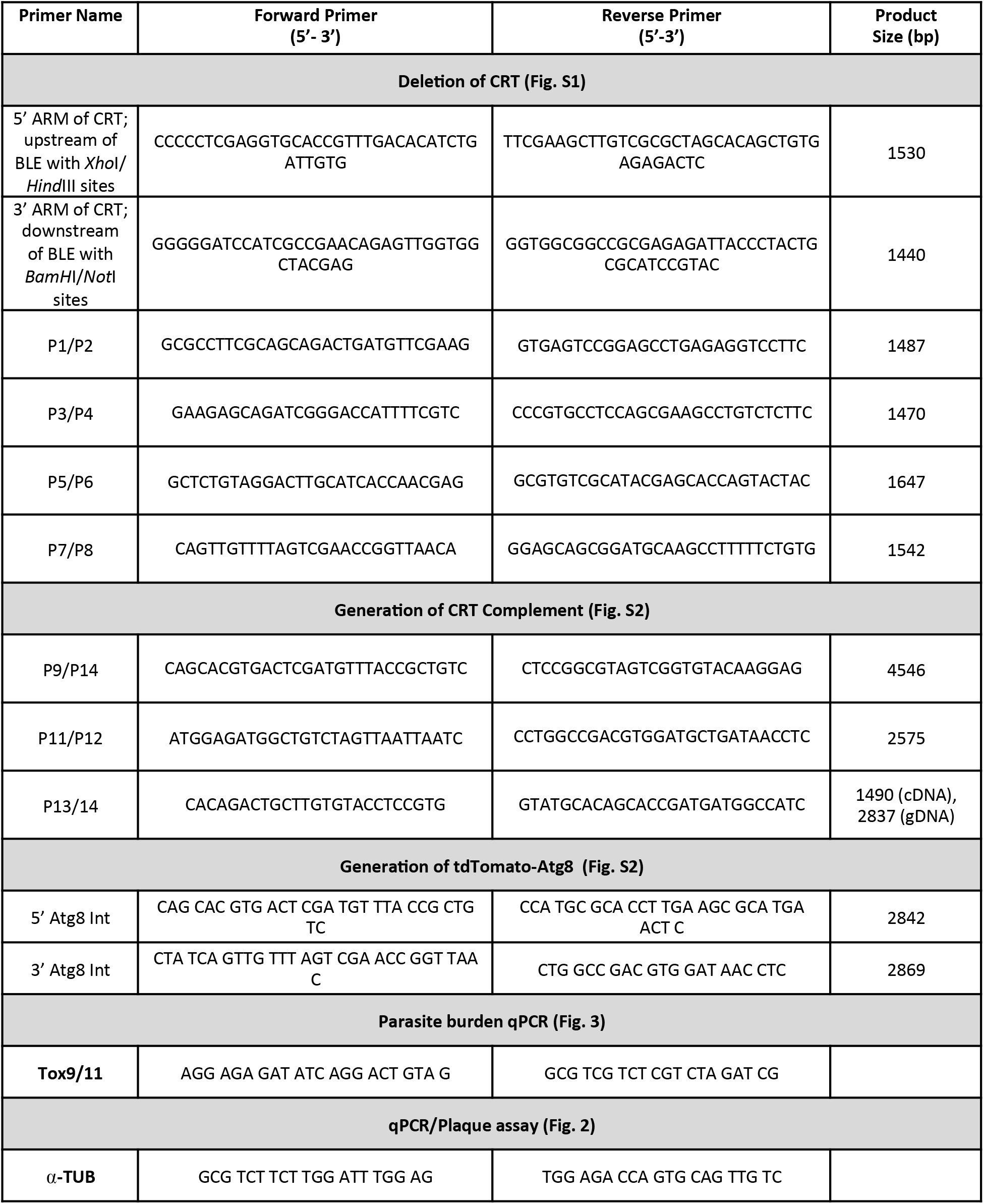
Primer sequences and PCR product sizes.

**Figure S1.**
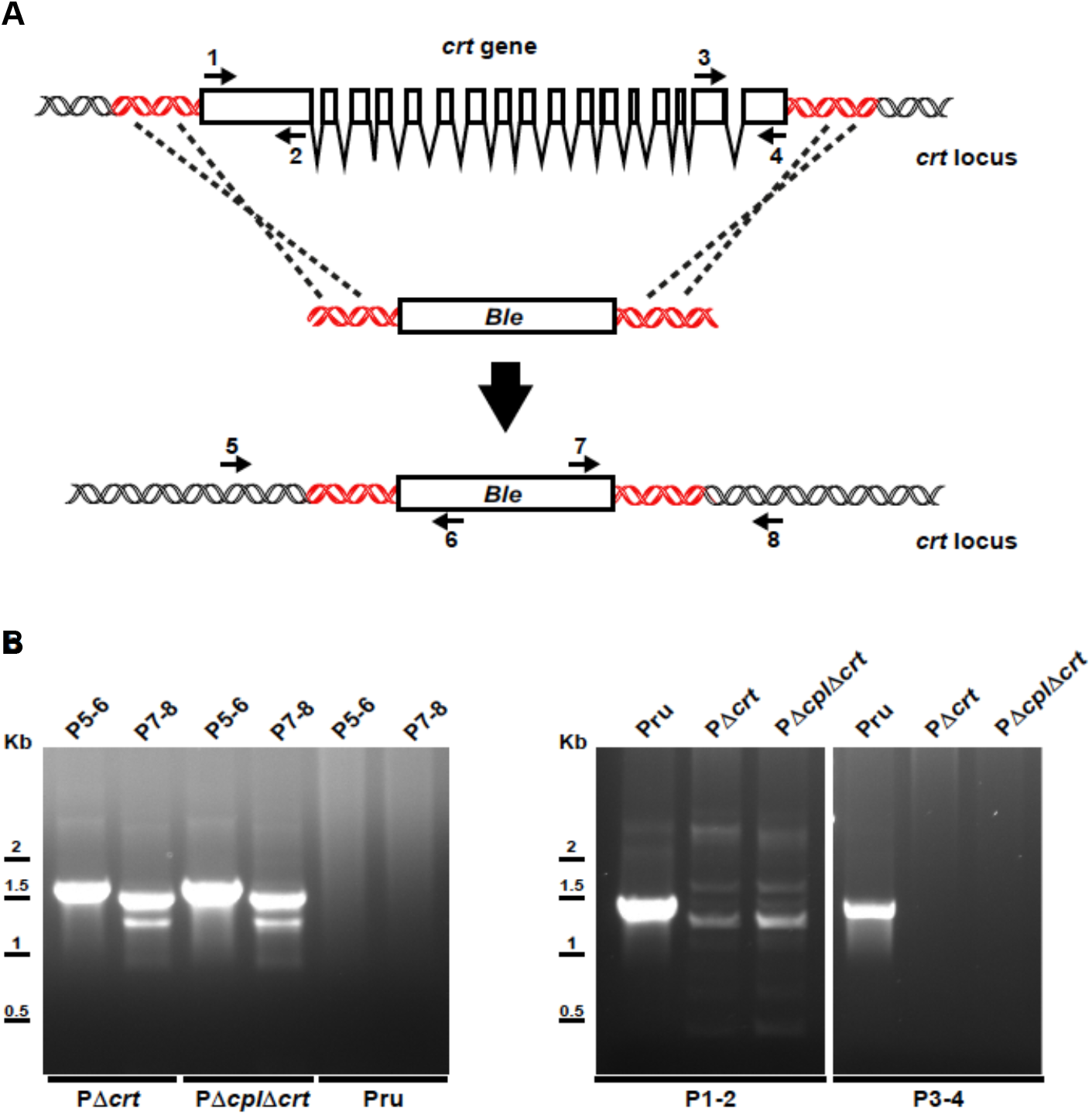
Targeted deletion of *CRT* in PRUΔku80 and PRUΔku80Δ*cpl*. A. A vector carrying the BLE selection cassette flanked at both ends by 1500 bp of homologous regions upstream and downstream of the *CRT* gene was used to delete *CRT* by double cross-over homologous recombination. B. Deletion of *CRT* was confirmed by PCR analyses using the primers indicated in each lane of the gel. Primer positions are shown in panel A. Primer sequences are provided in Table S1.

**Figure S2:**
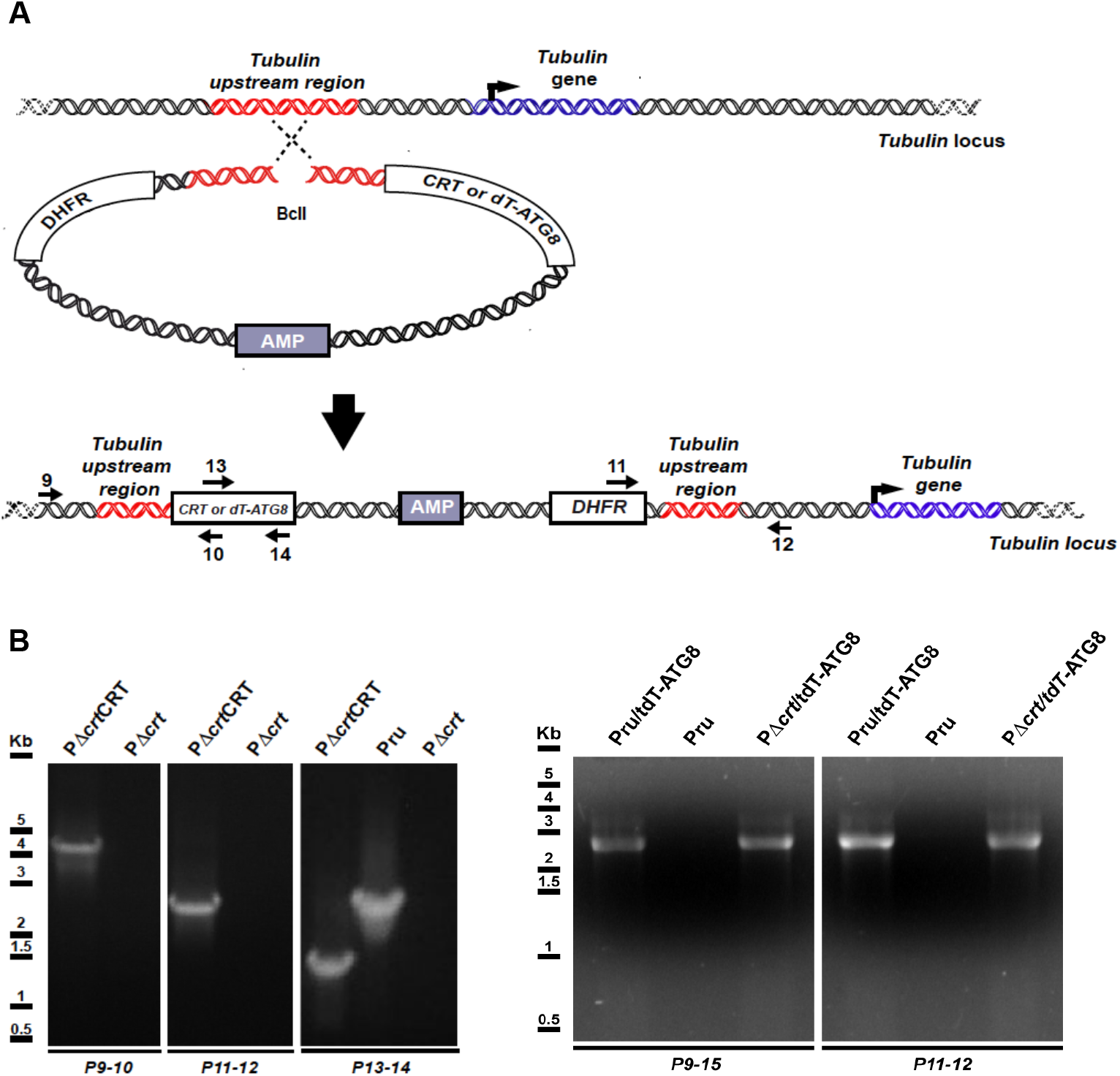
Genetic complementation of *CRT* and integration of dT-ATG8. A. Complementation of *CRT* was accomplished by integrating a plasmid carrying the *CRT* cDNA cloned downstream of 1000 bp of *CRT* 5’UTR to drive transcription of these sequences. The plasmid was integrated upstream of the tubulin gene by introducing in the complement plasmid a 1425 bp fragment encompassing this locus and linearization using the *BcII* to induce single cross-over. The tdTomato-Tg Atg8 expression cassette was integrated in the tubulin locus of the Δ*crt* strain using the same strategy described for the *CRT* complementation strain. B. Integration in the selected genome locus of the *CRT* complement or tdTomato-TgAtg8 plasmid was confirmed by PCR analysis. Primers used in these PCRs are indicated in panel A. Primer sequences are provided in Table S1.

**Figure S3:**
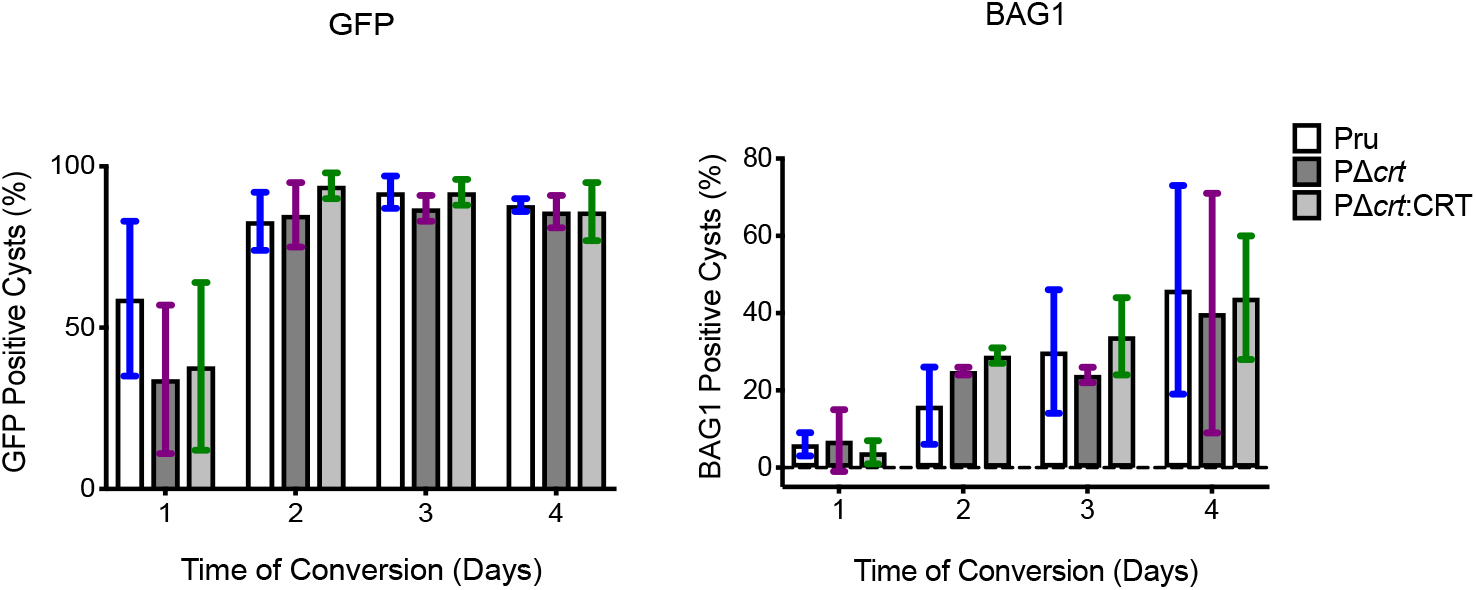
*In vitro* differentiation kinetics. Tachyzoite conversion into bradyzoite cysts was assessed over 4 days as determined by expression of GFP under the early bradyzoite promoter LDH2 or the late bradyzoite marker BAG1 by IF staining. Bars are presented as mean +/− S.D. of 3 independent experiments. The following parasitophorous vacuoles (PVs) were assessed on days 1, 2, 3, 4, respectively. Experiment 1: Pru (88, 175, 187, 144), PΔ*crt* (113, 166, 186, 204), PΔ*crt:CRT* (87, 124, 189, 182). Experiment 2: Pru (73, 84, 200, 248), PΔ*crt* (88, 137, 205, 190), PΔ*crt:CRT* (59, 101, 109, 117). Experiment 3: Pru (64, 90, 69, 97), PΔ*crt* (80, 63, 135, 214), PΔ*crt:CRT* (65, 78, 96, 132). One-way ANOVA with Tukey’s multiple comparisons was performed for comparing genotypes on each day.

**Figure S4.**
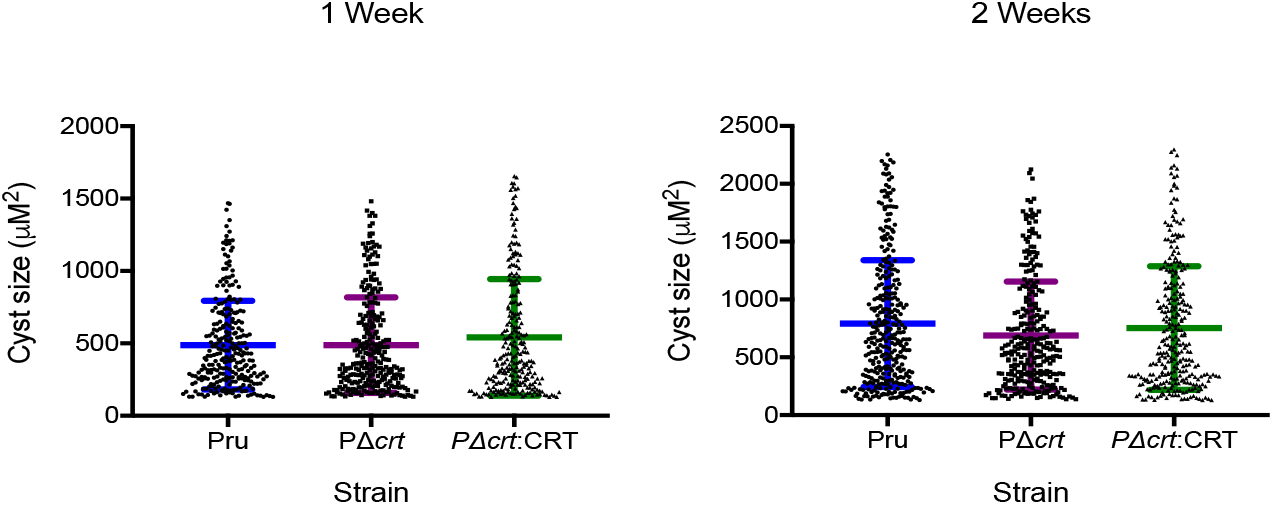
CRT deficiency does not alter *in vitro* cyst size. Cyst size after 1 and 2 weeks of *in vitro* differentiation. Line represents mean +/− S.D. of bradyzoite cysts in 3 independent experiments. The following number of cysts were analyzed for each experiment for week 1: Pru (78, 52, 142), PΔ*crt* (49, 90, 135), PΔ*crt:CRT* (89, 84, 87) and for week 2: Pru (111, 124, 102), PΔ*crt* (65, 131, 95), PΔ*crt:CRT* (90, 107, 102). Kruskal Wallis with Dunn’s multiple comparisons was performed.

**Figure S5.**
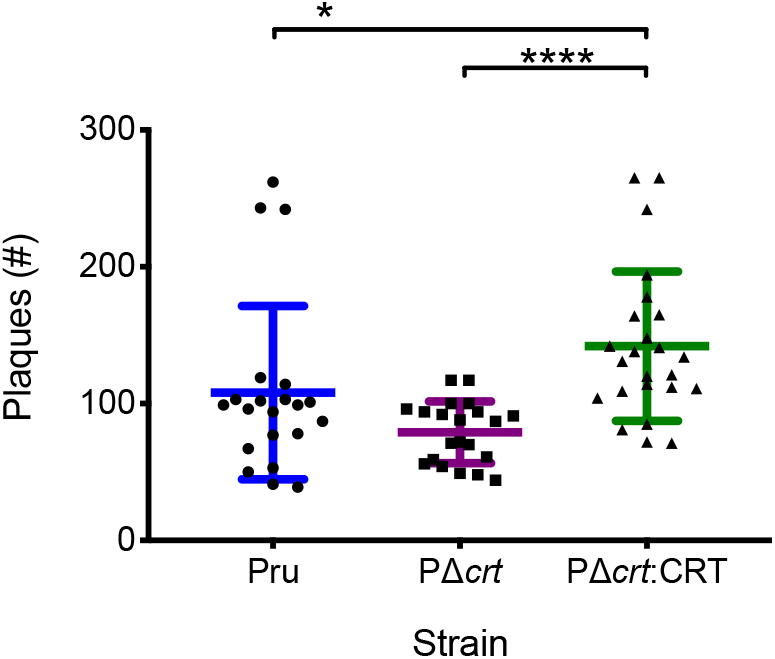

